# *C. elegans* Afadin is required for epidermal morphogenesis and functionally interfaces with the cadherin-catenin complex and RhoGAP PAC-1/ARHGAP21

**DOI:** 10.1101/2023.07.28.551013

**Authors:** Allison E. Hall, Diana Klompstra, Jeremy Nance

**Affiliations:** Department of Cell Biology, NYU School of Medicine, New York NY 10016; Skirball Institute of Biomolecular Medicine, NYU School of Medicine, New York NY 10016

## Abstract

During epithelial morphogenesis, the apical junctions connecting cells must remodel as cells change shape and make new connections with their neighbors. In the *C. elegans* embryo, new apical junctions form when epidermal cells migrate and seal with one another to encase the embryo in skin (‘ventral enclosure’), and junctions remodel when epidermal cells change shape to squeeze the embryo into a worm shape (‘elongation’). The junctional cadherin-catenin complex (CCC), which links epithelial cells to each other and to cortical actomyosin, is essential for *C. elegans* epidermal morphogenesis. RNAi genetic enhancement screens have identified several proteins that interact with the CCC to promote epidermal morphogenesis, including the scaffolding protein Afadin (AFD-1), whose depletion alone results in only minor morphogenesis defects. Here, by creating a null mutation in *afd-1*, we show that *afd-1* provides a significant contribution to ventral enclosure and elongation on its own. Unexpectedly, we find that *afd-1* mutant phenotypes are strongly modified by diet, revealing a previously unappreciated maternal nutritional input to morphogenesis. We identify functional interactions between AFD-1 and the CCC by demonstrating that E-cadherin is required for the polarized distribution of AFD-1 to cell contact sites in early embryos. Finally, we show that *afd-1* promotes the enrichment of polarity regulator and CCC-interacting protein PAC-1/ARHGAP21 to cell contact sites, and identify genetic interactions suggesting that *afd-1* and *pac-1* regulate epidermal morphogenesis at least in part through parallel mechanisms. Our findings reveal that *C. elegans* AFD-1 makes a significant contribution to epidermal morphogenesis and functionally interfaces with core and associated CCC proteins.

## Introduction

Epithelial cell shape changes play critical roles in the morphogenesis of organs and developing embryos. For example, coordinated shape changes within an epithelial cell sheet can lead to its elongation, thinning, or invagination (Honda, 2017; Lemke and Nelson, 2021; Leptin, 2005; Vuong-Brender et al., 2016). In addition, entire sheets of epithelial cells can undergo coordinated movements, such as during epiboly, when ectoderm cells migrate over the surface of embryos to properly position the future epidermis (Bruce and Heisenberg, 2020; Chisholm and Hardin, 2005; Honda, 2017; Leptin, 2005). These cell rearrangements and movements are executed by a complex interplay of proteins involved in cell polarization, adhesion, epithelial junction formation, and cytoskeletal remodeling.

The *C. elegans* epidermis provides a model system to investigate the mechanisms of cell shape change and movement and their contribution to morphogenesis. The epidermis first differentiates during the middle stages of embryogenesis, when cells on the dorsal and lateral surfaces of the embryo polarize along their apicobasal axis and assemble apical junctions, allowing them to function coordinately as an epithelial sheet. The initial phase of epidermal morphogenesis, called ventral enclosure, occurs when epidermal cells undergo an epiboly-like migration towards the ventral surface (Chisholm and Hardin, 2005; Vuong-Brender et al., 2016). At the completion of ventral enclosure, epidermal cells that were originally on opposite sides of the embryo meet at the ventral surface and seal together by forming new apical junctions with their contralateral partners (epidermal cells also undergo anterior movements that extend the epidermis anteriorly to the future oral cavity (Lynch and Hardin, 2009)). Following the completion of ventral enclosure, a subsequent phase of morphogenesis called elongation occurs when epidermal cells elongate along their anterior-posterior axis, which transforms the elliptical embryo into its final worm-like shape (Armenti and Nance, 2012; Chisholm and Hardin, 2005; Cram, 2014; Lynch and Hardin, 2009; Vuong-Brender et al., 2016). While elongation is initially driven by shape changes within epidermal cells, later stages of elongation require mechanical signaling inputs produced by contraction of underlying muscle cells (Armenti and Nance, 2012; Chisholm and Hardin, 2005; Cram, 2014; Lynch and Hardin, 2009; Vuong-Brender et al., 2016).

Proteins involved in the polarization, formation, and function of apical junctions are essential for the proper execution of ventral enclosure and elongation. Apical junctions in *C. elegans* epithelial cells contain two juxtaposed domains. The cadherin-catenin complex (CCC), which mediates cell adhesion and its linkage to the cytoskeleton, localizes most apically. Core components of the CCC include the transmembrane domain homophilic adhesion protein HMR-1/E-cadherin; HMP-2/β-catenin, which binds the E-cadherin cytoplasmic tail; and HMP-1/⍺-catenin, which binds to both β-catenin and F-actin (Costa et al., 1998; Lynch and Hardin, 2009). Mutations in any of the core CCC components disrupts the formation of new junctions during ventral enclosure, causing the embryo to rupture. DLG-1/Discs Large and its binding partner AJM-1 localize just basal to the CCC (Koppen et al., 2001; McMahon et al., 2001). DLG-1 and AJM-1 are required for junction maturation, and their loss causes embryos to arrest during elongation.

While the CCC plays a central role in junction formation, many additional proteins are required for remodeling and their loss often leads to less severe morphogenesis phenotypes or phenotypes only in compromised genetic backgrounds. Previously we showed that the conserved RhoGAP, PAC-1/ARHGAP21, colocalizes with CCC components at apical junctions in epidermal cells, where it acts to negatively regulate the activity of the RhoGTPase cytoskeletal regulator CDC-42 (Anderson et al., 2008). While mutations in *pac-1* do not disrupt epidermal morphogenesis significantly on their own, removing *pac-1* function in embryos containing a hypomorphic mutation in *hmp-1/⍺-catenin* greatly enhances epidermal morphogenesis defects and leads to near complete embryonic lethality (Zilberman et al., 2017). In non-epithelial cells of the early embryo, PAC-1 promotes cell polarity by localizing to cell contact sites, in part through catenin-mediated interactions with the HMR-1/E-cadherin cytoplasmic tail (Klompstra et al., 2015). One mechanism of interaction occurs through the linker protein PICC-1, which physically couples PAC-1 to the HMR-1/E-cadherin-binding catenin JAC-1/p120 (Klompstra et al., 2015). HMP-1/⍺-catenin also promotes PAC-1 localization, but how it does so is unknown. To further understand the connection between HMP-1/⍺-catenin and PAC-1, we investigated the role of the ⍺-catenin interacting protein Afadin/AFD-1, a large scaffolding protein that in other systems can bind to F-actin and has been shown to interact with ⍺-catenin.

Afadin has been most intensively studied in *Drosophila*, which has a single Afadin homolog named Canoe (Cno) (Choi et al., 2011; Jurgens et al., 1984; Miyamoto et al., 1995). Canoe promotes the initial stages of apical-basal polarity and helps link the actin cytoskeleton to junctions during apical constriction (Choi et al., 2013; Manning et al., 2019; Ooshio et al., 2007; Sawyer et al., 2009; Yu and Zallen, 2020). Cno also becomes enriched at tricellular junctions, where it is required for cell intercalation and recruiting E-cadherin to aid in cell adhesion (Yu and Zallen, 2020). Zygotic null Cno mutants are embryonic lethal, but have a range of mild to strong head involution defects (Manning et al., 2019; Sawyer et al., 2009). Mammalian Afadin is also an actin-binding protein that colocalizes with the CCC at adherens junctions (Mandai et al., 1997). Decreased Afadin function leads to an increase in tumorigenicity in mice and enhances invasion of breast cancer cells in culture (Chatterjee et al., 2012; Elloul et al., 2014; Fournier et al., 2011; Saito et al., 1998). Additionally, mammalian Afadin aids Par3 in the formation of adherens junctions (Ooshio et al., 2007).

The *C. elegans* Afadin homolog AFD-1 has been less extensively studied. AFD-1 immunoprecipitates with members of the CCC, and in intestinal cells and early embryos, colocalizes with HMR-1/E-cadherin (Callaci et al., 2015; Pickett et al., 2022; Slabodnick et al., 2023). Although *afd-1* loss-of-function mutations have not been described, inhibiting *afd-1* using RNAi enhances the embryonic arrest caused by the hypomorphic *hmp-1(fe4)* mutation in ⍺-catenin, and leads to minor defects in apical constriction during gastrulation (Callaci et al., 2015; Lynch et al., 2012; Slabodnick et al., 2023).

Here we characterize the expression, localization, and function of *afd-1* by modifying the endogenous *afd-1* locus. We show that *afd-1* mutants have significant defects in epidermal morphogenesis and AFD-1 colocalizes with the CCC within apical junctions of epidermal cells. Using the early embryo as a system to investigate how the CCC influences AFD-1, we find that AFD-1 is able to enrich at the plasma membrane independently of HMR-1/E-cadherin, but that HMR-1 promotes the polarized recruitment of AFD-1 to sites of cell-cell contact. Finally, we find that AFD-1 promotes PAC-1 localization, and we identify genetic interactions between *afd-1* and *pac-1* during epidermal morphogenesis. Our findings reveal new connections between the CCC, Afadin, and PAC-1 that contribute to robust epithelial morphogenesis.

## Materials and Methods

### WORM CULTURE AND STRAINS

All worms were maintained at room temperature 22°C (unless otherwise specified) on NGM plates seeded with *E. coli* OP50. Worms were maintained as previously described (Brenner, 1974). For nutritional analysis worms were either grown on NGM plates seeded with OP50, NGM plates seed with *E. coli* HT115, or RNAi plates containing IPTG seeded with HT115 transformed with pPD129.36 (empty RNAi vector) (Sturm et al., 2018; Timmons et al., 2001). All strains used in this paper are listed in the Key Resrouce Table.

### MICROSCOPY

#### Live imaging of embryos

Live imaging of embryos was performed on fresh 4% agarose pads in M9 buffer. All imaging with the exception of auxin and FRAP experiments was performed using a spinning disc confocal microscope (Nikon Eclipse Ti2, CSU-W1 spinning disk, 100X 1.35NA Silicone oil immersion objective, 488nm and 561nm lasers, matched Andor 888 Live EMCCD cameras). Live imaging of DLG-1^mCherry^ experiments were performed on the spinning disk confocal microscope with images taken every three minutes and 0.3 µm z stacks. FRAP and auxin experiments were imaged using a Leica SP8 confocal microscope (63X 1.4NA oil immersion objective, Argon tunable laser, and HyD detectors). DIC movies were acquired using an AxioImager (Zeiss), 63X 1.4 NA or 40× 1.3 NA objective, DIC optics, an Axiocam MRM camera, and AxioVision software. Timelapse images were acquired every 5 min for wild type and every 2 min for mutants for 7–9 h at 22°C, with *z*-planes spaced at 2 µm intervals.

#### FRAP imaging

L4 stage HMR-1^GFP-AID^; AFD-1^YFP^ worms were grown with or without auxin (see auxin methods) at 23°C for 24 hours prior to imaging. 2-cell embryos from hermaphrodites grown with or without auxin were mounted on 4% agarose pads and allowed to develop to the 4-cell stage, at which point they were subjected to FRAP at the ABa/ABp cell contact. FRAP experiments were carried out on a Leica SP8 confocal microscope (63X 1.4NA oil immersion objective, Argon tunable laser, and HyD detectors) using the FRAP module. Three pre-bleach images were taken at one second intervals. An ROI corresponding to a central region of the ABa/ABp cell contact was bleached at 100% 415 nm laser power with four consecutive pulses. Images were acquired every second for sixty seconds post-bleach. Area bleached was measured for recovery and analyzed in ImageJ (NIH).

#### Immunostaining

Embryos were obtained by chopping adult hermaphrodites in 7 µl of water on a coverslip, placing them on a poly-L-lysine-coated slide, freeze cracking on dry ice, and fixing initially for 20 minutes in methanol at -20°C followed by paraformaldehyde fix as described (Anderson et al., 2008). The following primary antibodies were used: mouse anti-HA 1:1,000 (clone 16B12; Covance), rabbit anti-HMR-1 1:10,000 (Klompstra et al., 2015), mouse anti-PSD-95 (recognizes DLG-1) 1:200 (Affinity BioReagents), rat anti-mCherry 1:100 (clone 5F8; Antibodies-online), rat anti-GFP 1:1,000 (Nacalai Tesque), rabbit anti-GFP 1:1,000 (Ab6556.25; AbCam). The following secondary antibodies were used: Cy3 anti-mouse IgG (subclasses 1+2a+2b+3) 1:250 (Jackson ImmunoResearch), Alexa Fluor 488 anti-rabbit IgG (H+L) 1:1,000 (Molecular Probes), Alexa Fluor 488 anti-rat IgG (Jackson ImmunoResearch), Cy5 anti-mouse IgG 1:200 (Jackson ImmunoResearch), Alexa Fluor 647 anti-rabbit 1:200 (Molecular Probes), Cy5 anti-rat IgG 1:200 (Jackson ImmunoResearch).

### IMAGE ANALYSIS

All images were analyzed using FIJI (ImageJ, NIH). For elongation rates, eggshell length was determined and used as a reference for one-fold elongation. The zero timepoint for all measurements was at the early comma stage (Zilberman et al., 2017).

#### Colocalization analysis

For colocalization of CCC members and AFD-1^YFP^ in epidermal cells, bean stage embryos were analyzed. A central plane was chosen for analysis, visualizing only the HMR-1 channel for images analyzing HMR-1, or only the DLG-1 or PAC-1 channel for figures 3D-E, so as not to bias the analysis. A plane where the most apical point of the epidermis beginning to enwrap the embryo was visible, with at least four distinct points of fluorescence in view was chosen. A 6 µm line was drawn starting from the most apical portion of the embryo (see Fig 3B for cartoon). Fluorescence intensity of both fluorophores was determined and averaged for two different junctions along the embryo. Data were normalized to account for differences in fluorescence intensity by setting the brightest point for each fluorophore to an arbitrary unit of 1. Zero was set as the most apical protein imaged. Graphs shown in figure 3 are the mean and standard deviation (cloud around the dark line) of the embryos. To determine significance, the distance between peaks was recorded and a two-tailed, paired t-test was performed.

#### Polarity index / HMR-1 independence experiments

To determine the polarity index of AFD-1^YFP^ localization in HMR-1^GFP-AID^ embryos, the YFP channel was analyzed in embryos from mothers grown with or without HMR-1 (with or without auxin). Contact fluorescence was measured as a 40-pixel line along the ABa/ABp contact of 4-cell embryos. Contact-free surfaces were measured as an average of the fluorescence intensity of a line trace along the cortex of the ABa cell. Contact enrichment was calculated as the contact intensity divided by two to account for the double membrane, and this value was divided by the intensity at the contact free surface. Significance was determine using an unpaired, two-tailed, t-test.

Contact vs. contact-free fluorescence intensity was measured as a 6 µm line drawn from the outside of the embryo across the ABa cortex (for contact-free measurement) or from ABa into ABp (for contact measurement). Five lines were measured across the membranes and the image background was subtracted from every point along the line. An average of the five lines for both membranes was taken and the brightest point along the line was set as zero. Data were normalized to set the brightest point (the membrane) to one in the +HMR-1 (-auxin) embryos. The -HMR-1 (+auxin) fluorescence intensity was normalized to these data, allowing us to determine the relationship between AFD-1^YFP^ fluorescence intensity at the contact-free and contact membranes with and without HMR-1. To determine significance, the peak fluorescence intensity for each embryo with and without auxin was compared and an unpaired, two-tailed, t-test was used.

#### Embryonic elongation analysis

Elongation rates were measured as previously described (Zilberman et al., 2017). The length of the eggshell was determined and used as a reference for one-fold elongation. The zero-time point for all measurements was at the comma stage.

#### PAC-1^GFP^ contact enrichment analysis

*pie-1p::pac-1::gfp* and *pie-1p::pac-1::gfp*; *afd-1(Δ)* embryos were placed on slides together to ensure equal imaging parameters between strains. A single z-plane at the center of 4-cell embryos was taken. For contact enrichment analysis, fluorescence intensity along a 40-pixel line on the ABa/ABp cell contact was taken and divided by the average of four individual lines within the cytoplasm of ABa (ensuring no line crossed the nucleus). The image background was subtracted from both the contact and average cytoplasm values before contact fluorescence was divided by cytoplasm fluorescence.

#### FRAP analysis

Pre-bleach and post-bleach images were concatenated using ImageJ (NIH). The fluorescence recovery of AFD-1^YFP^ was measured and recorded using the Plot Z-axis profile function. FRAP data were normalized, as described (Day et al., 2012), to pre-bleach levels at the ABa/ABp contact and accounting for photobleaching over time by measuring the ABa/EMS contact for comparison with image background subtracted. These data were plotted over the sixty second imaging course for +/-HMR-1 (-/+ auxin). The time to half recovery was calculated as the point in time on the recovery curve in which half of the final fluorescence had been reached, calculated by: ((t_final_ – t_i_)/2)+t_i_ (t_i_ being the first bleach image taken). Data are from three independent experiments and significance was calculated using a two-tailed, unpaired, t-test.

### CRISPR GENOME ENGINEERING

#### afd-1::yfp

The plasmid for CRISPR/Cas9 genome editing to make *afd-1(xn54: afd-1-yfp + unc-119)* was constructed as described previously (Dickinson et al., 2013), with the following modifications. The guide RNA sequence from plasmid pDD122 was replaced with the sequences (5′-TCCTGGCTTTTGCGGGGTGG - 3′) to create a single guide RNA (sgRNA) that cleaves near the *afd-1* C-terminus (plasmid pDK50). A homologous repair plasmid for *afd-1* (pDK49) was constructed using Gibson assembly. The following DNA segments were assembled in order: 1501 bp upstream of *afd-1* stop codon (including one silent point mutation to destroy the PAM site) as the left homology arm; *yfp* with *unc-119*; and the 3′-terminal 1502 bp of *afd-1* genomic sequence as the right homology arm. *yfp* with *unc-119* flanked by LoxP sites was amplified from plasmid pJN601, which contains LoxP-flanked *unc-119* inserted in reverse orientation into a synthetic intron within *yfp* (Armenti et al., 2014). The vector backbone for each construct was PCR-amplified from pJN601 using Gibson assembly primers that overlapped with homology arms for *afd-1*.

#### Other *afd-1* modifications

All other strains were made using CRISPR/Cas9-mediated genome editing as previously described (Paix et al., 2017). Cas9 (Berkeley) was pre-incubated with crRNA and tracrRNA (IDT), mixed with a ssDNA oligonucleotide (IDT) containing ∼35bp homology arms, and injected into worms using the *dpy-10(cn64)* co-CRISPR repair strategy (Paix et al., 2017). F1 Roller worms were singled and genotyped by PCR to identify edited worms. crRNA, repair template sequences and genotyping primers are listed in the Key Resources Table.

#### afd-1(Δ)

The *afd-1(Δ)* mutant was created using the STOP-IN method of introducing an early stop codon (Wang et al., 2018). We modified this protocol to exclude the exogenous Cas9 target site and replaced that with 3 X HA tags just prior to the early stop codon in order to determine if a truncated protein was produced. The ssDNA repair oligo was injected into *afd-1::yfp/tmC20* balanced worms (Dejima et al., 2018). Once heterozygous mutant worms were identified, they were rebalanced with a clean *tmC20* balancer from an outcross.

#### afd-1::3XHA

A ssDNA oligonucleotide containing three HA tags inserted just before the *afd-1* stop codon was used as a repair template. CRISPR/Cas9-mediated editing was used as described above, injecting into a wild type (N2) strain.

#### afd-1(xn195)

A ssDNA oligonucleotide altering the splice acceptor site just 5’ of leucine 1278 from a G to an A was used to create *afd-1(xn195)*. The ssDNA oligonucleotide also introduced three silent mutations within exon 18 (isoform a, WormBase) to prevent crRNA recutting. This was injected into *afd-1::yfp* (FT1586).

#### afd-1::yfp^reverted^

Two crRNAs were used to cut out the three HA tags inserted to make *afd-1(Δ)*, one targeting the first of the three HA tags and the other just 3’ of the inserted STOP-IN cassette. A ssDNA oligonucleotide that restored wild-type *afd-1* sequence was used and injected into *afd-1(Δ)*.

### AUXIN EXPERIMENTS

Auxin plates were made as NGM plates with a final concentration of 2mM auxin. Auxin indole-3-acetic acid (IAA) was purchased from Alfa Aesar (#A10556). A 200 mM stock of auxin was made in ethanol and diluted 1:100 in cooled NGM media. Auxin stocks were made fresh, and control plates contained equivalent amounts of ethanol in NGM. For auxin treatment, L4 *hmr-1::ZF1::gfp::aid*; *afd-1::yfp*; *sun-1p::TIR-1* worms were placed on plates with and without auxin at 23° C for 24 hours. Embryos from treated mothers were placed on 4% agarose pads and imaged. To ensure there was consistency in imaging, imaging was paired with +/-auxin treated embryos on the same slide.

### EMBRYONIC ARREST

Seven to ten L4 worms were placed on a seeded NGM plate at midday and allowed to develop and lay eggs at 23°C until the following morning, when mothers were transferred to a new plate and allowed to lay eggs for 8 hours before transferring once more. Eggs were counted upon removal of the mother and unhatched embryos were counted 24 hours later. Experiments were performed a minimum of three times.

### ISOLATION OF *afd-1* SPLICE SITE MUTATION

A synchronized population of *afd-1::yfp* L4 hermaphrodites was mutagenized with EMS as previously described (Brenner, 1974) and allowed to self for two generations. F3 embryos were scored in utero for AFD-1^YFP^ cell contact localization using an AxioImager (Zeiss), 63X 1.4 NA or 40× 1.3 NA objective. Mutants were recovered directly from slides. *afd-1(xn195)* was created by introducing the identified G to A splice acceptor site mutation into a clean AFD-1^YFP^ background using CRISPR/Cas9 genome editing.

### STATISTICAL ANALYSIS

Statistical analysis was performed using GraphPad Prism 9. For elongation experiments, unpaired t-tests were used to compare length at the final 60 minute time point. For embryonic arrest experiments, a Fisher’s exact test was used. For all other data, unpaired, two-tailed t-tests were used. The data in graphs are presented using SuperPlots (Lord et al., 2020). For SuperPlots, individual data from at least three independent experiments are color-coded and represented as small, transparent dot; the mean for each experiment is shown as a larger dot with corresponding color, and the error bars represent the standard error of the mean. Statistical test, sample size, and p-values are reported in the figure legends.

## Results

### *afd-1* mutants show diet-influenced embryonic arrest

Prior studies utilizing RNAi to inhibit *afd-1* function revealed that *afd-1* knockdown strongly enhanced the lethality of a hypomorphic *hmp-1/α-catenin* mutant, but only caused a low level of embryonic lethality in an otherwise wild-type genetic background (Lynch et al., 2012; Serre et al., 2023; Slabodnick et al., 2023). To determine if disruption of the *afd-1* gene results in a more severe developmental phenotype, we modified the *afd-1* locus by first inserting *yfp* immediately prior to the stop codon [*afd-1(xn54)*; hereafter *afd-1::yfp,* encoding AFD-1^YFP^], then introducing an early stop codon (STOP-IN cassette) preceded by three HA epitope tags [*afd-1(xn124)*; hereafter *afd-1(*ϕλ)] (Fig. 1A). We chose the 15^th^ exon of *afd-1* isoform a for the stop cassette because it is the most 5’ exon common to all identified *afd-1* isoforms and splice variants (Wormbase version WS288). *afd-1(*ϕλ) is predicted to encode a truncated product lacking an intact PDZ domain as well as the C-terminal half of the protein (Fig. 1A).

**Figure 1.**
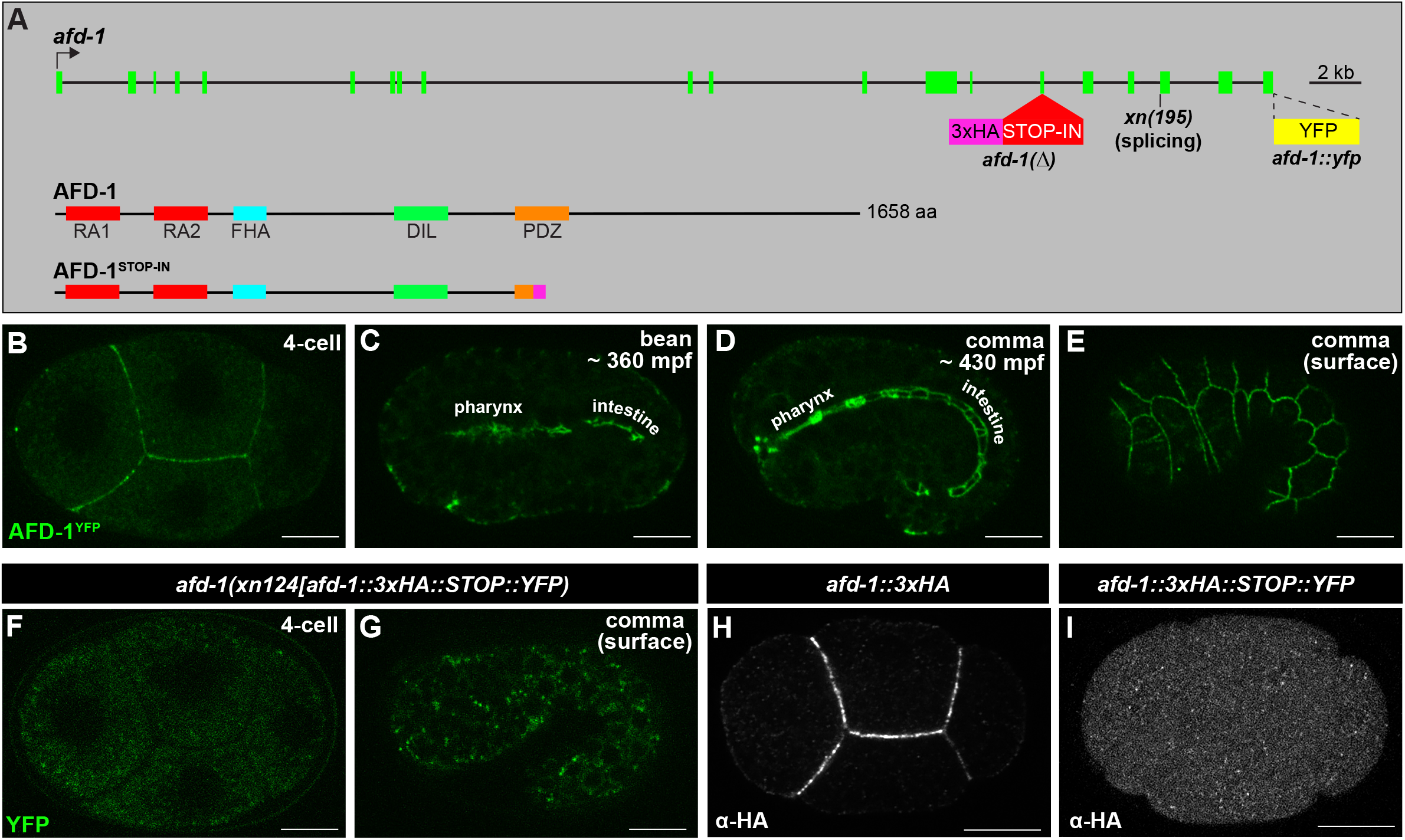
AFD-1 localizes to cell junctions. **(A)** Above: schematic of *afd-1::yfp*, *afd-1(Δ)*, and *afd-1(xn195)*. *afd-1(Δ)* introduces 3XHA tags 5’ of a STOP-IN cassette. Below: schematic of AFD-1 protein with predicted truncation caused by the STOP-IN cassette. Domains: RA (Ras-associated), FHA (Fork-head associated), DIL (Dilute domain). **(B-E)** Live imaging of AFD-1^YFP^ localization to cell contacts in the 4-cell embryo (B) and junctions in the bean (C) and comma (D-E) stage embryos, showing both internal and surface views. Developing pharynx and intestine are identified. **(F-G)** Live imaging of *afd-1(Δ)* in the 4-cell (F) and comma stage (G) embryo, showing a surface view. **(H-I)** Immunostaining against HA tags in endogenously tagged *afd-1::3xHA* (H) and *afd-1(Δ)* (I). Scale bars, 10 µm.

In agreement with previous studies that examined AFD-1 localization (Lynch et al., 2012; Pickett et al., 2022; Serre et al., 2023; Slabodnick et al., 2023), we observed AFD-1^YFP^ at cell contacts in early embryonic cells of *afd-1::yfp* embryos (Fig. 1B) and at apical junctions in both internal epithelial cells (Fig. 1C-D; bean stage ∼360 m.p.f., comma stage ∼430 m.p.f.) and epidermal cells (Fig. 1E), which form during mid-embryogenesis. Consistent with the introduced stop codon affecting all *afd-1* mRNAs, we did not detect AFD-1^YFP^ expression in *afd-1(*11) embryos (Fig. 1F-G). *afd-1(*11) mutant embryos did not show HA immunoreactivity above background levels (Fig. 1I), in contrast to embryos from an *afd-1::3xha* strain [*afd-1(xn214)*] (Fig. 1H), suggesting either that the early stop codon induces nonsense-mediated decay of the *afd-1* mRNA (Arribere et al., 2020) or that the truncated protein produced is unstable. These observations indicate that the *afd-1(*11) allele is likely to be a null mutation.

*afd-1(*11) homozygous mutants arose from self-fertilized heterozygous mothers at the expected frequency (154/649 embryos, 24%) and grew to the adult stage. However, 26% of the progeny of *afd-1(*11) mutant mothers (*afd-1^M-Z-^* embryos) arrested during embryonic development (Table 1), in contrast to the minimal lethality reported previously in *afd-1(RNAi)* embryos (Lynch et al., 2012; Serre et al., 2023). Unexpectedly, in the course of performing RNAi experiments on *afd-1* mutants, we observed that the embryonic arrest of *afd-1(*11) mutants was strongly influenced by the *E. coli* strain provided as a food source to their *afd-1(*11) mutant mothers: in side-by-side feeding experiments, we observed 23% lethality when *afd-1(*11) mutant mothers were fed OP50 bacteria, but only 9% lethality when worms were grown on plates seeded with the HT115 strain used in RNAi experiments (Table 2). Suppression of *afd-1(*11)-induced lethality by HT115 was not related to RNAi, because HT115 alone suppressed the lethality of *afd-1(*11) embryos to a similar extent as HT115 bacteria transformed with a control RNAi vector (L4440) and seeded on RNAi-inducing plates (containing IPTG) (Table 2). Although OP50 and HT115 are both *E. coli* strains, their metabolomic profiles show some significant differences (Neve et al., 2020; Reinke et al., 2010), suggesting that one or more *E. coli* metabolites modulates an *afd-1*-mediated essential developmental event. These findings could also explain why *afd-1(*11) mutants reared on OP50 bacteria have more severe embryonic lethality than that reported for *afd-1(RNAi)* embryos, which were grown on HT115 (Lynch et al., 2012; Serre et al., 2023).

**Table 1.** Embryonic lethality of *afd-1* mutants.

| Genotype<sup>#</sup> | Embryonic lethality | *n* | Significance (p) |
| --- | --- | --- | --- |
| wild type | 1.5% | 2164 | N/A |
| *afd-1*(Δ) | 26% | 5111 | <0.0001<sup>&</sup> |
| *afd-1::yfp* | 12% | 818 | N/A |
| *afd-1::yfp<sup>reverted</sup>* | 17% | 3078 | <0.001<sup>*</sup> |

<sup>#</sup>Maternal and zygotic genotype
<sup>&</sup>Compared to wild type, Fisher's exact test
<sup>\*</sup>Compared to *afd-1::yfp*, Fisher's exact test

**Table 2.** Nutrient dependence of embryonic lethality in *afd-1* mutants.

| Genotype<sup>#</sup> (food source) | Embryonic lethality | *n* | Significance (p)<sup>&</sup> |
| --- | --- | --- | --- |
| *afd-1*(Δ) (*E. coli* OP50) | 23% | 2682 | N/A |
| *afd-1*(Δ) (*E. coli* HT115) | 9% | 1768 | <0.0001 |
| *afd-1*(Δ) (*E. coli* HT115 + L4440 vector) | 12% | 936 | <0.0001 |

<sup>#</sup>Maternal and zygotic genotype
<sup>&</sup>Compared to *afd-1*(Δ) (*E. coli* OP50), Fisher's exact test

### *afd-1* is required for epidermal morphogenesis but not epidermal junction formation

To determine the cause of embryonic lethality in *afd-1(*11) embryos, we recorded time-lapse differential interference contrast (DIC) movies of their development. Of 24 *afd-1(*Δ) embryos imaged, 12/24 arrested prior to elongation due to epidermal ruptures at the ventral surface or at the embryo anterior. The other 12/24 completed ventral enclosure and began to elongate, but within this class, 4/12 arrested before completing elongation (8/8 wild-type embryos completed elongation). Embryos that were able to elongate did so more slowly than wild type (Figure 2A-C, Movies 1-2). The higher degree of embryonic arrest in our DIC time-lapse movies compared to *afd-1(*11) embryos laid on plates is not unexpected given that imaged embryos are compressed beneath a coverslip.

**Figure 2.**
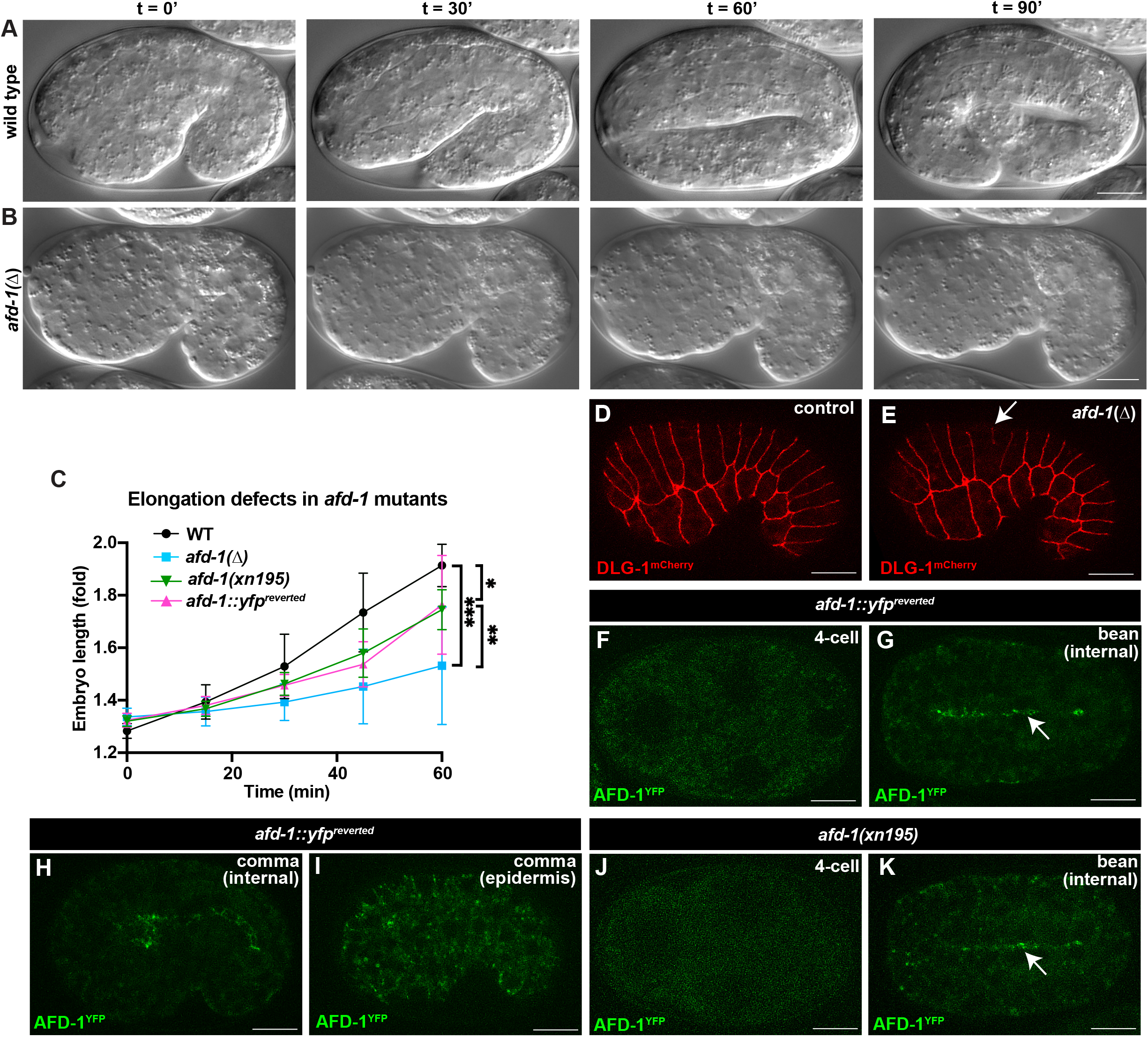
*afd-1* mutations cause elongation defects. **(A-B)** Stills from DIC movies of wild type (A) and *afd-1(Δ)* (B) at indicated times. **(C)** Elongation rates measured from DIC movies of wild type n = 8, *afd-1(Δ)* n = 12, *afd-1(xn195)* n= 12, and *afd-1::yfp^reverted^* worms n = 6. Data plotted are mean values with standard deviation as error bars. Significance determined for the final time point (60 min) using an unpaired t-test. t = 0’ represents the comma stage. ***p ≤ 0.001, **p ≤ 0.01, *p ≤ 0.05. **(D-E)** Maximum projection of four z-stacks from live imaging of comma stage embryos from control (DLG-1^mCherry^) and *afd-1(Δ)* (*afd-1(Δ)*; DLG-1^mCherry^) showing epidermal DLG-1^mCherry^. Arrow in E indicates fusing dorsal epidermal cells. **(F-I)** Maximum projection of five stacks from live imaging of *afd-1::yfp^reverted^*at the 4-cell (F) bean (G), and comma stages (H-I) Arrow indicates developing intestinal junctions (G). Internal view (H) and surface view of the epidermis (I) of a comma stage embryo. **(J-K)** Maximum projection of five stacks from live imaging of *afd-1(xn195)* embryos as the 4-cell (J) and bean (K) stage. Arrow indicates developing intestine. Scale bars, 10 µm.

Because mutations in genes required for epidermal junction formation also cause embryonic arrest prior to or during elongation (e.g. *par-6, pkc-3/apkc, dlg-1/discs large, let-413/scribble*) (Bossinger et al., 2001; Firestein and Rongo, 2001; Koppen et al., 2001; Legouis et al., 2000; McMahon et al., 2001; Montoyo-Rosario et al., 2020; Totong et al., 2007), we examined epidermal junctions in *afd-1(*Δ) embryos in a strain expressing the endogenously tagged junction component DLG-1^mCherry^ (Kroll et al., 2021). *afd-1(*Δ) embryos imaged in the *dlg-1::mCherry* background showed a similar suite of epidermal phenotypes. However, unlike mutations that affect junction formation, in which DLG-1 puncta never assemble into continuous apical belts, DLG-1^mCherry^ formed continuous belts around epidermal cells in *afd-1(*Δ) mutants (24/24 *dlg-1::mCherry* embryos, 66/66 *afd-1(*Δ); *dlg-1::mCherry* embryos) (Fig. 2D-E). Together, these observations indicate that *afd-1* is important for epidermal morphogenesis but is dispensable for forming the apical junctions that connect epidermal cells together.

To confirm that the phenotypes we observed in *afd-1(*Δ) mutants were caused by the introduced premature stop codon, we used CRISPR to revert *afd-1(*Δ) back to *afd-1::yfp* [*afd-1(xn202)*; hereafter *afd-1::yfp^reverted^*]. Surprisingly, doing so restored zygotic AFD-1^YFP^ expression in the epithelial cells of mid-stage embryos (Fig. 2G), but not maternal AFD-1^YFP^ expression in early embryos (Fig. 2F). We speculate that the maternal silencing of *afd-1::yfp^reverted^* is a result of RNAe – a piRNA-mediated germline silencing mechanism that targets mRNAs containing foreign sequences, like *yfp* (Shirayama et al., 2012). *afd-1::yfp^reverted^*embryos showed a significantly lower level of embryonic lethality than *afd-1(*Δ) mutant embryos (17% versus 26%) but significantly higher than *afd-1::yfp* alone (12%) (Table 1), and mutant embryos elongated at a rate intermediate between that of wild type and *afd-1(Δ)* (Fig. 2C). Levels of AFD-1^YFP^ in epithelial cells and epidermal cells of *afd-1::yfp^reverted^* embryos appeared markedly lower than AFD-1^YFP^ in *afd-1::yfp* embryos (Fig. 2H,I; compare to Fig. 1C-E), potentially explaining the intermediate phenotype of *afd-1::yfp^reverted^*embryos.

In the course of a genetic screen to identify mutants that affect AFD-1^YFP^ localization in early embryos (see Methods), we identified a mutant, *afd-1(xn195)*, with a remarkably similar phenotype to that of *afd-1::yfp^reverted^*. In *afd-1(xn195)* embryos, visible maternal AFD-1^YFP^ expression was absent (Fig. 2J), and zygotic expression of AFD-1^YFP^ in epidermal cells appeared substantially reduced (Fig. 2K). *afd-1(xn195)* embryos also showed an intermediate rate of elongation very similar to that of *afd-1::yfp^reverted^* embryos (Fig. 2C). The causative mutation in *afd-1(xn195)* (see Methods) is a splice acceptor mutation three exons from the 3’ end of the gene (Fig. 1A). Because both *afd-1::yfp^reverted^* and *afd-1(xn195)* block maternal expression of AFD-1^YFP^ and cause a similar phenotype, we speculate that the *afd-1(xn195)* splice site mutation affects a maternal *afd-1* splice variant. Together, the phenotypes of *afd-1::yfp^reverted^*and *afd-1(xn195)* indicate that maternal *afd-1* expression contributes to epidermal morphogenetic events that occur during mid-embryogenesis.

### Afadin colocalizes with components of the cadherin-catenin complex in the epidermis

Given the epidermal morphogenesis defects we observed in *afd-1* null mutants, we examined AFD-1^YFP^ subcellular localization in epidermal cells relative to other proteins known to regulate epidermal morphogenesis. HMR-1/E-cadherin (together with associated catenins HMP-1/⍺-catenin and HMP-2/β-catenin) is required for ventral enclosure and localizes to apical junctions connecting epidermal cells (in addition to apical junctions in other epithelia). DLG-1/Discs large, which is dispensable for ventral enclosure but essential for elongation (Bossinger et al., 2001; Firestein and Rongo, 2001; Koppen et al., 2001; McMahon et al., 2001), also localizes to apical junctions but in a domain just basal to HMR-1/E-cadherin; when viewed using confocal microscopy, the two proteins show partially overlapping distributions with distinct locations of peak intensity (Koppen et al., 2001; McMahon et al., 2001) (Fig 3A-B). In immunostained embryos, AFD-1^YFP^ colocalized with HMR-1/E-cadherin but not DLG-1 (Fig 3C-D’), consistent with previous findings in *C. elegans* intestinal cells and studies from other systems showing that Afadin and ⍺-catenin can interact (Callaci et al., 2015; Pickett et al., 2022). The RhoGAP PAC-1/ARHGAP21 also localizes to apical junctions and regulates epidermal elongation redundantly with HMP-1/⍺-catenin (Zilberman et al., 2017). As with HMR-1/E-cadherin, AFD-1^YFP^ colocalized with PAC-1 at epidermal apical junctions (Figure 3E).

**Figure 3.**
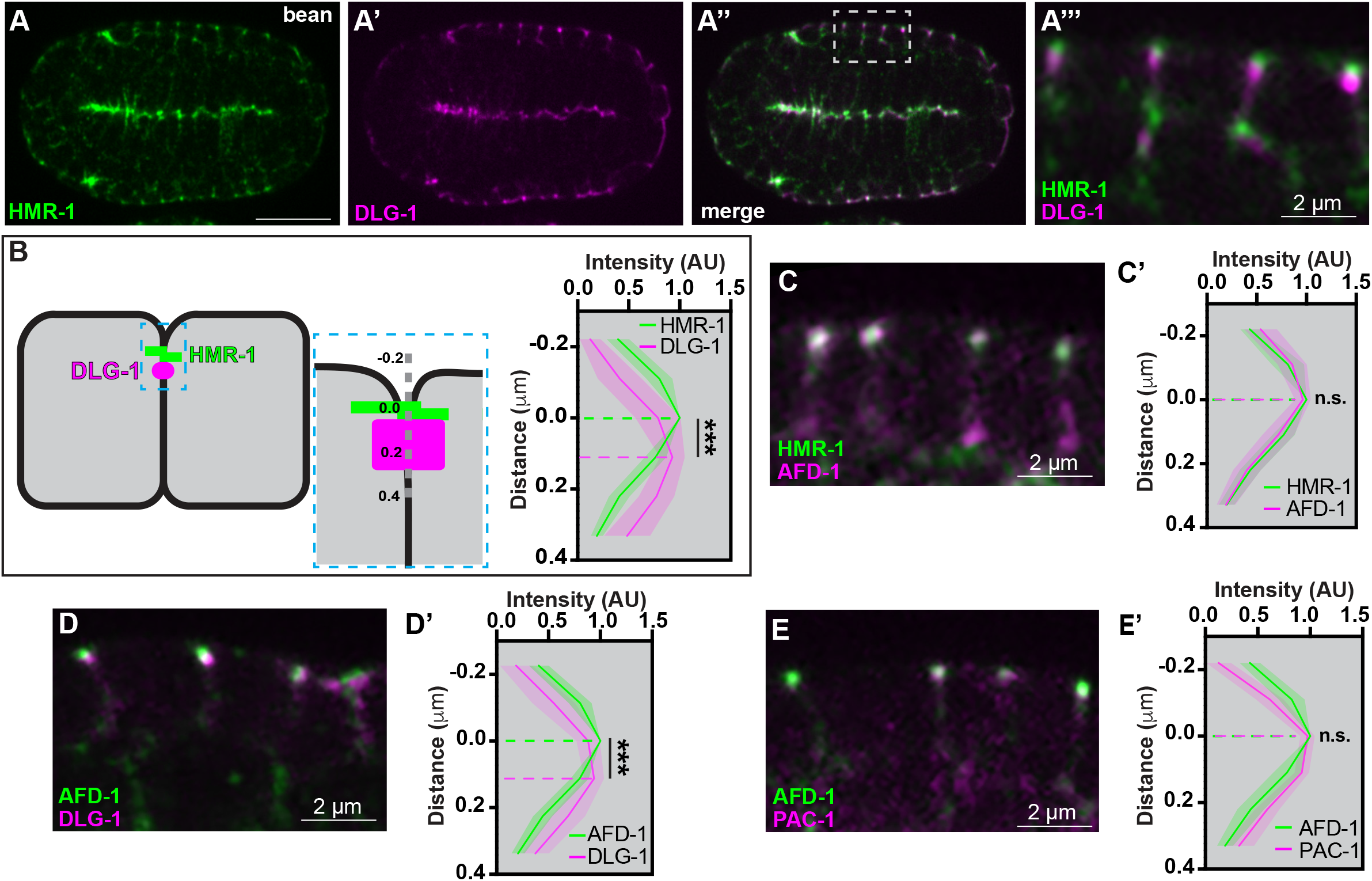
AFD-1 colocalizes with the cadherin-catenin complex in the developing epidermis. **(A-A’’’)** Staining of HMR-1 and DLG-1 in wild type bean stage embryos. A’’’ shows 500% zoom of the dashed box indicated region in A’’. **(B)** Left: schematic of colocalization analysis used in A and all subsequent images. A zoom in of the blue dashed box indicates the line drawn from apical to basal within epithelia and the resulting graph (right) showing arbitrary fluorescence units and HMR-1 peak centered on 0. Graph displays the mean (dark line) of n = 12 embryos with standard deviation shown as a surrounding cloud of lighter color. Significance calculated using a paired t-test on the distance of the peak intensity from the most apical point along the drawn line. **(C-C’)** Representative image and resulting analysis of staining of HMR-1 and AFD-1^YFP^ in *afd-1::yfp* embryos. 500% zoom of the epidermis shown (C) and analysis of n = 17 embryos. **(D-D’)** Representative image and resulting analysis of staining of AFD-1^YFP^ and DLG-1 in *afd-1::yfp* embryos. 500% zoom of the epidermis shown (D) and analysis of n = 12 embryos. **(E-E’)** Representative image and resulting analysis of staining of AFD-1^YFP^ and PAC-1 ^mCherry^ in *afd-1::yfp*; *pac-1::mCherry* embryos. 500% zoom of the epidermis shown (E) and analysis of n = 15 embryos. In all graphs peak intensities for each fluorophore were normalized to a value of 1. Dashed lines indicate peak fluorescence intensity. Graphs display the mean intensity (dark line) with standard deviation (cloud around the line). Paired t-tests were used to determine significance ***p ≤ 0.001, n.s. p > 0.05. Scale bar, 10 µm except where indicated.

### HMR-1/E-cadherin promotes AFD-1 cell contact enrichment

Given the colocalization of AFD-1^YFP^ and HMR-1/E-cadherin, we asked if *hmr-1* regulates AFD-1 localization. We initially examined early embryos, where *hmr-1* is dispensable for cell adhesion and mutant embryos appear morphologically normal. AFD-1^YFP^ and HMR-1/E-cadherin both localize to cell contacts between early embryonic cells. Because *hmr-1* is difficult to inhibit completely using RNAi (Chihara and Nance, 2012), we depleted HMR-1/E-cadherin protein using the auxin inducible degron (AID) system. For these experiments, we combined an endogenously tagged *hmr-1::gfp::aid* allele (Lee et al., 2021) with a germline TIR1 driver (*sun-1*p*::TIR1*); TIR1 targets AID-tagged proteins for degradation only if auxin is present (Nishimura et al., 2009; Sepers et al., 2022; Zhang et al., 2015). Without auxin, when HMR-1^GFP-AID^ was present, AFD-1^YFP^ enriched strongly at cell-cell contacts (Fig 4A,A’). We quantified AFD-1^YFP^ contact enrichment by calculating the ‘polarity index’, which we defined as half of the average intensity at a specific cell contact divided by the full average intensity at a contact-free surface of the same cell (Fig. 4C). This calculation adjusts for the presence of two apposing plasma membranes at the cell-cell contact site; proteins that are uniformly enriched at all cell surfaces should have a polarity index of 1, and proteins enriched at contact sites should have a polarity index significantly greater than 1. In control embryos where auxin was absent and HMR-1^GFP-AID^ was present, the polarity index of AFD-1^YFP^ was ∼2 (Fig. 4C). However, when auxin was present and HMR-1^GFP-AID^ was depleted, the polarity index of AFD-1^YFP^ was reduced to ∼1, similar to the polarity index of a GFP-fused PH_PLC1∂1_ domain that is thought to localize uniformly to the plasma membrane (Audhya et al., 2005) (Fig 4B-D). This finding is consistent with a previous report showing a decrease in N-terminally tagged AFD-1^mKate2^ at cell contacts in *hmr-1(RNAi)* early embryos, although this study did not examine AFD-1^mKate2^ levels at contact-free surfaces (Slabodnick et al., 2023). When compared directly, AFD-1^YFP^ signal at contact-free surfaces was significantly higher when HMR-1^GFP-AID^ was depleted, and conversely, AFD-1^YFP^ signal was significantly lower at cell contacts when HMR-1^GFP-AID^ was depleted compared to no auxin controls (Fig 4E). Consistent with HMR-1/E-cadherin stabilizing AFD-1 at cell contacts, in FRAP experiments, the time to half recovery (ι−_½_) of AFD-1^YFP^ photobleached at a specific cell contact was modestly but significantly shorter when HMR-1^GFP-AID^ was depleted, suggesting that without HMR-1, AFD-1 moves more freely in the membrane (Fig 4F-G). Because *hmr-1* mutants develop severe epidermal morphogenesis defects, and HMR-1/E-cadherin contributes to junction formation (Costa et al., 1998; Pickett et al., 2022; Raich et al., 1999), quantitative comparisons of AFD-1^YFP^ in wild-type and *hmr-1* mutant epidermal cells are difficult to interpret. However, we noted that at least some AFD-1^YFP^ was still able to enrich at apical junctions in epidermal cells in *hmr-1* mutant embryos (Fig 4H-I’). Taken together, our observations in early embryos and epithelial cells show that AFD-1 is able to localize in the absence of HMR-1/E-cadherin, but at least in the early embryo, HMR-1/E-cadherin helps concentrate AFD-1 at sites of cell contact where the two proteins colocalize.

**Figure 4.**
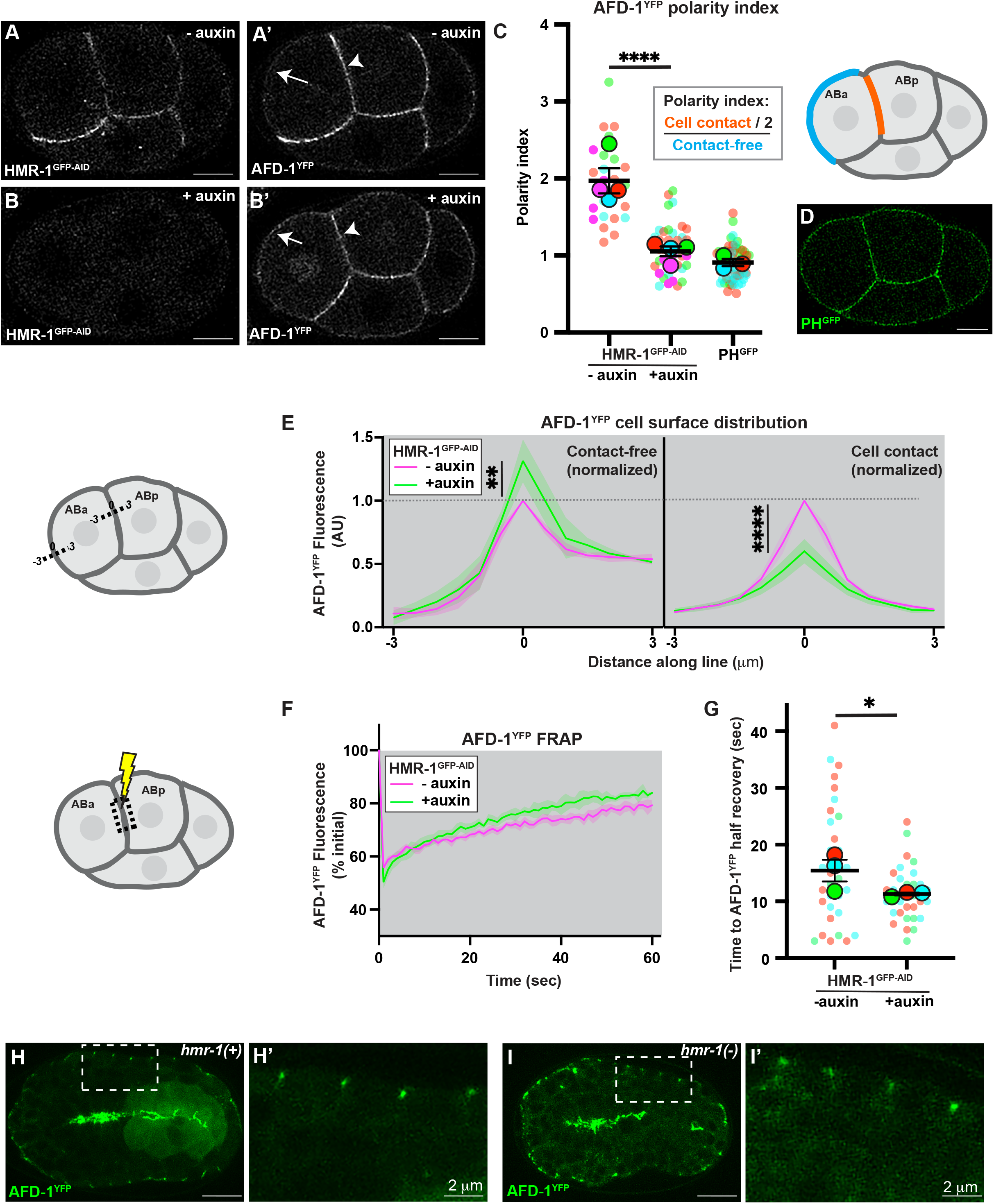
HMR-1 promotes AFD-1 cell contact enrichment. **(A-B’)** Live images of a 4-cell *hmr-1::gfp::aid; afd-1::yfp* embryo from a mother expressing germline specific TIR-1 without auxin (A-A’) and with auxin (B-B’). Arrowheads indicate contact enrichment of AFD-1^YFP^ and arrows indicate further enrichment at non-contacts when auxin is present. **(C)** Quantification of polarity index of AFD-1^YFP^. Polarity index was measured as shown in the cartoon and formula. **(D)** Representative image of a 4-cell embryo expressing PH^GFP^. **(E)** AFD-1^YFP^ fluorescence at contact-free (left) and cell contacts (right). Cartoon depicts the method of analysis. All data from three independent experiments were normalized relative to the no auxin AFD-1^YFP^ membrane peak, set to a value of 1. Dark pink lines indicate the mean fluorescence without auxin, standard deviation is shown as the lighter cloud around the line. Dark green lines indicate the mean fluorescence with auxin, standard deviation shown as the lighter cloud around the line. Unpaired t-tests were used to determine significance between peak intensities, n > 22 embryos for each condition **p ≤ 0.01, ****p ≤ 0.0001. **(F)** AFD-1^YFP^ FRAP recovery with and without auxin in *hmr-1::gfp::aid; afd-1::yfp* embryos. Cartoon depicts region of bleach. Graph displays t = 0 as the pre-bleach image and fluorescence recovery over sixty seconds. Dark lines indicate the mean recovery of three independent experiments, n = 33 embryos for no auxin and n = 34 embryos with auxin; standard deviation shown as a lighter cloud around the line. **(G)** Time to half recovery of the data presented in (F). **(H-I’)** Maximum projection of 5 z stacks of live embryos showing AFD-1^YFP^ localization in bean stage embryos in *hmr-1* mutants with and without a *hmr-1(+)* rescuing array. Images in H’ and I’ are 500% zoom in of the epidermis from the dashed box in H and I. The presence of the *hmr-1(+)* rescuing array is indicated by the green in the center of the embryo in (H) (expression from an *end-1p::gfp* cotransformation marker). Scatter dot plots: individual data points from three or four independent color-coded experiments are shown as transparent dots, the mean of each experiment is shown as the larger dot of corresponding color, the mean of the means is shown as a black line and S.E.M is indicated by the error bars. Unpaired t-test was used to determine significance, *p<0.05, ****p ≤ 0.0001. Scale bar, 10 µm except where indicated.

### AFD-1 promotes PAC-1 contact enrichment in the early embryo

The biochemical function of AFD-1 in regulating morphogenesis is unclear. Given that AFD-1^YFP^ colocalizes with HMR-1/E-cadherin and PAC-1/ARHGAP21, together with previously reported findings that *afd-1(RNAi)* and a *pac-1* mutant strongly enhance the lethality and epidermal defects of a hypomorphic *hmp-1/⍺-catenin* mutant (Lynch et al., 2012; Zilberman et al., 2017), we examined the genetic relationship of *afd-1* with *hmp-1* and *pac-1*. We first attempted to confirm that the *afd-1(Δ)* mutant, like *afd-1(RNAi)*, enhances the hypomorphic *hmp-1(fe4)* mutant (which causes ∼50% lethality on its own). Although we were able to generate strains that were homozygous for one mutant and heterozygous for the other *(hmp-1; afd-1(Δ)/Balancer* and *afd-1(Δ); hmp-1/Balancer*), neither strain produced any viable double homozygotes (0/105 from *afd-1(Δ); hmp-1/Balancer,* and 0/200 from *hmp-1; afd-1(Δ)/Balancer*), consistent with strong genetic interactions between the two genes. Next, we examined genetic interactions between *afd-1* and *pac-1*, which on its own shows a lower level of lethality than *afd-1(Δ)* (Table 3). Two different mutant alleles of *pac-1* (*xn6,* which introduces a stop codon prior to the RhoGAP domain; and *xn1,* which disrupts a splice site) each significantly enhanced the lethality of *afd-1(Δ)*. Because the *pac-1(xn6)* mutation is predicted to be a functional null (Anderson et al., 2008), these experiments suggest that *afd-1* and *pac-1* both regulate epidermal morphogenesis, but do so at least in part through separate mechanisms rather than via a simple linear pathway.

**Table 3.** Genetic interactions between *afd-1* and *pac-1*.

| Genotype | Embryonic lethality | *n* | Significance (p) |
| --- | --- | --- | --- |
| *afd-1*(Δ)<sup>*</sup> | 26% | 5111 | N/A |
| *pac-1*(*xn6*) | 10% | 2580 | N/A |
| *afd-1*(Δ); *pac-1*(*xn6*) | 39% | 796 | <0.0001<sup>&</sup> |
| *pac-1*(*xn1*) | 7% | 1802 | N/A |
| *afd-1*(Δ); *pac-1*(*xn1*) | 42% | 834 | <0.0001<sup>&</sup> |

<sup>\*</sup>Data from Table 1
<sup>&</sup>Compared to *afd-1*(Δ), Fisher's exact test

Finally, to determine if AFD-1 and PAC-1 functions intersect, we asked if *afd-1*, like *hmp-1* (Klompstra et al., 2015), regulates PAC-1 localization. We performed these experiments in early embryos, where AFD-1^YFP^ and PAC-1^GFP^ are both enriched at cell contacts. Measuring levels of PAC-1^GFP^ both at cell contacts and within the cytoplasm, we observed a significant decrease in PAC-1^GFP^ contact-enrichment (average level at a cell contact divided by average cytoplasmic level) in *afd-1(Δ)* mutant embryos, although very low levels of PAC-1^GFP^ remained at cell contacts (Fig 5A-C). We could not perform this analysis quantitatively in epidermal cells because PAC-1 can only be detected at this stage by immunostaining, although we noted that at least some PAC-1^mCherry^ detected by immunostaining was still present at apical junctions (Fig 5D-E). These observations, taken together with the genetic interaction experiments, suggest that while AFD-1 and PAC-1 must regulate epidermal morphogenesis through at least partially different mechanisms, one role for AFD-1 is promoting PAC-1 localization.

**Figure 5.**
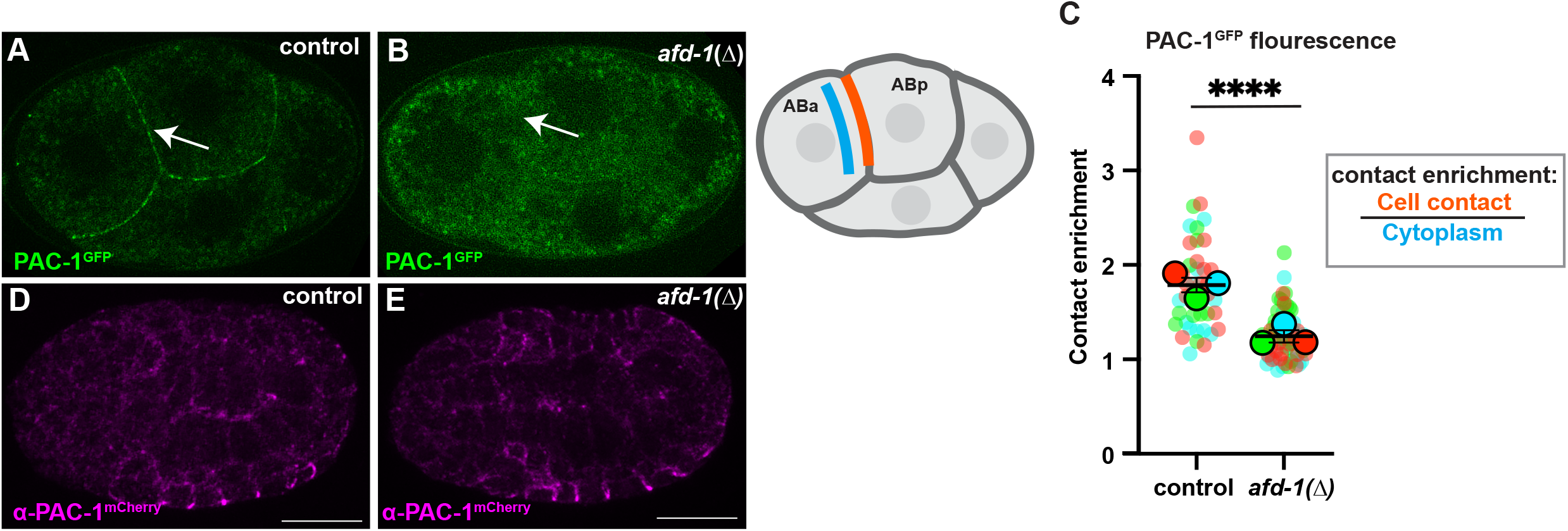
AFD-1 promotes PAC-1 contact enrichment in the early embryo. **(A-B)** Live imaging of 4-cell embryos expressing PAC-1^GFP^ in control or *afd-1* mutant embryos. Arrows indicate the ABa/ABp cell contact analyzed. Images were placed on a black background for figure continuity. **(C)** Contact enrichment quantification of data in (A) and (B). The cartoon depicts the method of analysis. Individual data points from three independent color-coded experiments are shown as transparent dots, the mean of each experiment is shown as the larger dot of corresponding color, the mean of the means is shown as a black line and S.E.M is indicated by the error bars. An unpaired t-test was used to determine significance ****p<0.0001. **(D-E)** Staining of PAC-1^mCherry^ in control and *afd-1(Δ)* bean stage embryos. Scale bar, 10 µm.

## Discussion

Our work describes a role for *C. elegans afd-1/Afadin* in epidermal morphogenesis. By generating the first described *afd-1* loss-of-function mutant, we showed that *afd-1* is required for both the completion of ventral enclosure and embryo elongation, which is driven in part by epidermal cell shape changes. A commonality between these two processes is that epidermal apical junctions must be created and remodeled as cells make new physical connections and alter their shapes. These mutant phenotypes are consistent with previously published observations showing that *afd-1(RNAi)* enhances the epidermal morphogenesis defects of a hypomorphic mutation in *hmp-1/⍺-catenin* (Lynch et al., 2012), and expand on these observations by showing that, in many embryos, *afd-1* is required for these events on its own. In both mammalian and fly systems Afadin has been shown to link to the actin cytoskeleton (Mandai et al., 1997; Sawyer et al., 2009), like ⍺-catenin. While a direct interaction between *C. elegans* AFD-1 and the actin cytoskeleton has not been demonstrated, several studies have now shown indirect connections (Lynch et al., 2012; Slabodnick et al., 2023). AFD-1 may therefore act alongside the CCC to help link junctions to the cytoskeleton during morphogenesis.

### AFD-1 localization and HMR-1/E-cadherin

Endogenously tagged AFD-1^YFP^ colocalizes with HMR-1/E-cadherin at cell contacts in early embryos and at apical junctions in epidermal cells, consistent with previous findings. AFD-1^YFP^ in the early embryo largely lost its contact enrichment when we depleted HMR-1 protein, consistent with findings made using N-terminally tagged AFD-1^mKate2^ and *hmr-1(RNAi)*. We expand on these results to show that AFD-1^YFP^ is not only reduced at cell contacts when HMR-1/E-cadherin is depleted, but that it also appears at contact-free surfaces, suggesting that HMR-1/E-cadherin is not required for AFD-1 localization but rather its polarity at the plasma membrane. We propose that AFD-1 itself, either through direct lipid binding or association with other proteins, enriches at the plasma membrane, and that HMR-1/E-cadherin helps to trap AFD-1 at cell contacts. This interpretation is supported by our FRAP experiments, which revealed that AFD-1^YFP^ photobleached at cell contacts is less mobile when HMR-1/E-cadherin is present than when it is depleted. In epidermal cells, AFD-1^YFP^ was still able to concentrate at apical junctions in *hmr-1* mutant embryos, suggesting that its localization in epithelial cells likely involves redundant mechanisms. The adhesion protein SAX-7 and the MAGUK MAGI-1 were previously shown in antibody staining experiments to promote AFD-1 localization to epidermal junctions (Lynch et al., 2012). Our findings in the early embryo raise the possibility that they may do so together with HMR-1/E-cadherin.

### Connection between AFD-1 and PAC-1

Previous work from our lab revealed a connection between the RhoGAP PAC-1/ARHGAP21, the RhoGTPase CDC-42, and F-actin dynamics at apical junctions in epidermal cells (Zilberman et al., 2017). In *pac-1* mutants, levels of HMR-1/E-cadherin and two of its interacting catenins, JAC-1/p120 catenin and HMP-1/⍺-catenin, increase in epidermal cell apical junctions. *pac-1* mutants also enhance the epidermal morphogenesis defects of hypomorphic *hmp-1(fe4)/a-catenin* mutants (Zilberman et al., 2017). Given our finding that PAC-1 and AFD-1 colocalize both at cell contacts in early embryos and at apical junctions in epidermal cells, one possibility is that these proteins work together to promote epidermal morphogenesis, at least in part through CDC-42 and F-actin regulation. Our observations suggest a more complex interplay between AFD-1 and PAC-1. In the early embryo, we showed that *afd-1* promotes PAC-1 localization to cell contact sites, providing a functional link between the two proteins. However, in epidermal cells, PAC-1 was still able to localize in *afd-1* mutants. Moreover, the embryonic lethality of *pac-1; afd-1* mutants was significantly higher than that of either single mutant. These observations suggest that AFD-1 and PAC-1 can work together, but do not do so exclusively. This interpretation is consistent with our previous findings in the early embryo that PAC-1 localization is regulated redundantly by separate catenin pathways *(hmp-1/⍺-catenin* and *jac-1/p120 catenin).* Given that Afadin can physically interact with ⍺-catenin, it is possible that AFD-1 interfaces with PAC-1 together with HMP-1/⍺-catenin, and that both PAC-1 and AFD-1 have additional targets important for morphogenesis. This functional relationship could be conserved, as work in mammalian cell culture using BioID showed an *in vivo* interaction between Afadin (AFDN) and the mammalian PAC-1 homolog, ARHGAP21 (Goudreault et al., 2022).

### Maternal and embryonic sources of *afd-1* and alternative splicing

Our mutant screen for factors required to localize *afd-1* yielded an interesting mutation in the *afd-1* gene itself. *afd-1(xn195)*, which mutates a splice acceptor upstream of *afd-1* exon 18 in the *afd-1::yfp* knock-in allele, specifically prevents expression of maternal AFD-1^YFP^. The maternal-specific phenotype of the *xn195* splicing mutation was likely explained when we found that reverting a non-sense mutation we introduced in *afd-1::yfp* caused a similar specific loss of maternal AFD-1^YFP^. The phenotype of *afd-1::yfp^reverted^* must be an epigenetic loss of maternal *afd-1::yfp* expression (likely through the RNAe silencing pathway, which targets foreign mRNA sequences), since the sequence itself is the same as *afd-1::yfp* (Shirayama et al., 2012). Putting these observations together, we infer that the *afd-1(xn195)* mutation affects a maternal splice form of *afd-1*, and that zygotically expressed *afd-1* can use an alternative splice variant that does not require the splice acceptor mutated in *afd-1(xn195)*. A large-scale analysis of exon usage by RNA-seq in *C. elegans* revealed that splicing using the canonical form of exon 18 occurs ∼90% of the time, while in ∼9% of cases the upstream exon is spliced to the middle of exon 18 through use of an alternative splice acceptor (Tourasse et al., 2017). This minor product with a shorter exon 18 would bypass the *afd-1(xn195)* mutation (which would be skipped and spliced out). Although we do not know whether the maternal and zygotic forms of *afd-1* have similar or distinct functions, the intermediate phenotype of *afd-1(xn195)* and *afd-1::yfp^reverted^* with respect to lethality and embryo elongation indicate that maternal *afd-1* products significantly contribute to epidermal morphogenesis within the embryo.

### The *afd-1* mutant phenotype is affected by diet

We were surprised to find that the embryonic lethality of *afd-1* mutants was highly dependent on the strain of *E. coli* fed to their mothers: *afd-1* mutants grown on OP50 showed 2.5 times the embryonic lethality as those grown on HT115. Although HT115 is used for RNAi experiments, we found that HT115 suppressed *afd-1* mutant lethality even when the bacteria lacked the plasmid required for producing dsRNAs, thereby uncoupling the suppression from RNAi. These findings strongly suggest that metabolic differences between OP50 and HT115 significantly modify the *afd-1* mutant phenotype. It is not without precedent that diet can affect RNAi phenotype. RNAi against a nuclear hormone receptor, *nhr-114*, when worms are grown on OP50 show >80% sterility, however this drops to <5% sterility when worms are grown on HT115 (Gracida and Eckmann, 2013). The cause of the sterility appears to be a decrease in the germline stem cell pool of the worms (Gracida and Eckmann, 2013). While this cannot be the case for *afd-1(Δ)* worms, it does indicate that diet can affect phenotype on various food sources. One possibility is that diet in the mother affects the embryonic levels of one or more proteins that function with AFD-1, such as CCC components or PAC-1.

### Author contributions

AEH designed, executed, and interpreted all experiments except for creation of *afd-1::yfp*, which was made by DK. JFN contributed to experimental design and data analysis. AEH and JFN wrote and edited the manuscript, and DK provided comments on the manuscript.

## Acknowledgements

We thank Mike Boxem (Utrecht University) and Kang Shen (Stanford University) for providing worm strains. Some strains were obtained from the *Caenorhabditis* Genetics Center (CGC), which is supported by the NIH Office of Research Infrastructure Programs (P40OD010440). We thank the members of the Nance laboratory and Dan McIntyre for comments on the manuscript. Microscopy used instrumentation in the NYULMC Microscopy Laboratory, which is partially supported by the Laura and Isaac Perlmutter Cancer Center support grant P30CA016087 from the National Institutes of Health/National Cancer Institute. This work was supported by a fellowship from the National Institutes of Health (NIH) (F32HD101227) to AEH, and a research grant from the NIH to JN (R35GM118081).

---

**Movie 1**. Elongation in a wild type embryo. Movie from the stills in Fig. 2A. DIC imaging, time points are every 5 min, display rate 7 frames/s.

**Movie 2**. Elongation in an *afd-1(Δ)* embryo. Movie from the still in Fig. 2B. DIC imaging, time points are every 2 min, display rate 7 frames/s.

## KEY RESOURCE TABLE

### Bacterial and Virus strains

KEY RESOURCE TABLE Bacterial and Virus strains
| Reagent or resource | Source | Identifier |
| --- | --- | --- |
| OP50, *E. coli* | *Caenorhabditis* Genetics Center | OP50 |
| RNAi feeding, *E. coli* | *Caenorhabditis* Genetics Center | HT115 |

### Chemicals, Peptides, and Recombinant Proteins

| Reagent or resource | Source | Identifier |
| --- | --- | --- |
| Cas9 protein | Univ. California Berkeley |  |
| Indole-3-acetic acid, 98+% | Alfa Aesar | Cat # A10556-14 |

### Experimental Models: Organisms/Strains

| Reagent or resource | Source | Identifier |
| --- | --- | --- |
| N2, wild type *C. elegans* | *Caenorhabditis* Genetics Center | N2 |
| *afd-1(xn54 [afd-1::yfp, unc-119(+)] I; unc-119(ed3) III* | This study | FT1586 |
| *afd-1(xn124 [afd-1::3xHA:STOP-IN::yfp) / tmC20 [unc-14(tmls1219) dpy-5(tm9715 myo-2P::Venus)] I* | This study | FT2205 |
| *afd-1(xn195) I; unc-119(ed3) III* | This study | FT2372 |
| *afd-1(xn202 [afd-1::yfp reverted]); unc-119(ed3) I* | This study | FT2384 |
| *afd-1(xn212[afd-1::3xHA])* | This study | FT2483 |
| *dlg-1(mib23[dlg-1::mCherry]) X* | Kroll et al. 2021 | BOX235 |
| *afd-1(xn124)/tmC20 I; dlg-1(mib23[dlg-1::mCherry-LoxP]) X* | This study | FT2411 |
| *afd-1(xn54) I; xnls178 [mCherry::pac-1, unc-119(+)] unc-119(ed3) III* | This study | FT2335 |
| *afd-1(xn124)/tmC20 I; xnls178 III* | This study | FT2517 |
| *hmr-1(wy1341[hmr-1::ZF1::gfp::AID\*]) afd-1(xn54) I; ieSi38 [sun-1p::TIR1::mRuby::sun1 3'UTR, unc-119(+)] IV* | This study | FT2467 |
| *Itls38 [pie-1p::GFP::PH<sub>PLC101</sub>, unc-119(+)]; zuls251 [nmy-2p::pgl-1::mRFP-1]; unc-119(ed3) III* | This study | FT554 |
| *hmr-1(zu389) I; xnEx42 [pW02-21(hmr-1), pdpy-7::RFP, pJN151(end-1::GFP), pBSKS+]; afd-1(xn54) I* | This study | FT2259 |
| *pac-1(xn6) unc-119(ed3) III; xnls27 [pie-1p::gfp::pac-1, unc-119(+)]* | Anderson et al. 2008 | FT99 |
| *xnls27; afd-1(xn124)/tmC20* | This study | FT2393 |
| *pac-1(xn6) III* | Anderson et al. 2008 | FT93 |
| *afd-1(xn124)/tmC20 I; pac-1(xn6) III* | This study | FT2332 |
| *pac-1(xn1) III* | Anderson et al. 2008 | FT45 |
| *afd-1(xn124)/tmC20 I; pac-1(xn1) III* | This study | FT2330 |
| *hmp-1(fe4); afd-1(xn124)/tmC20* | This study | FT2340 |
| *afd-1(xn124); hmp-1(fe4)/tmc12* | This study | FT2341 |

### ssDNA Oligonucleotides and RNAs

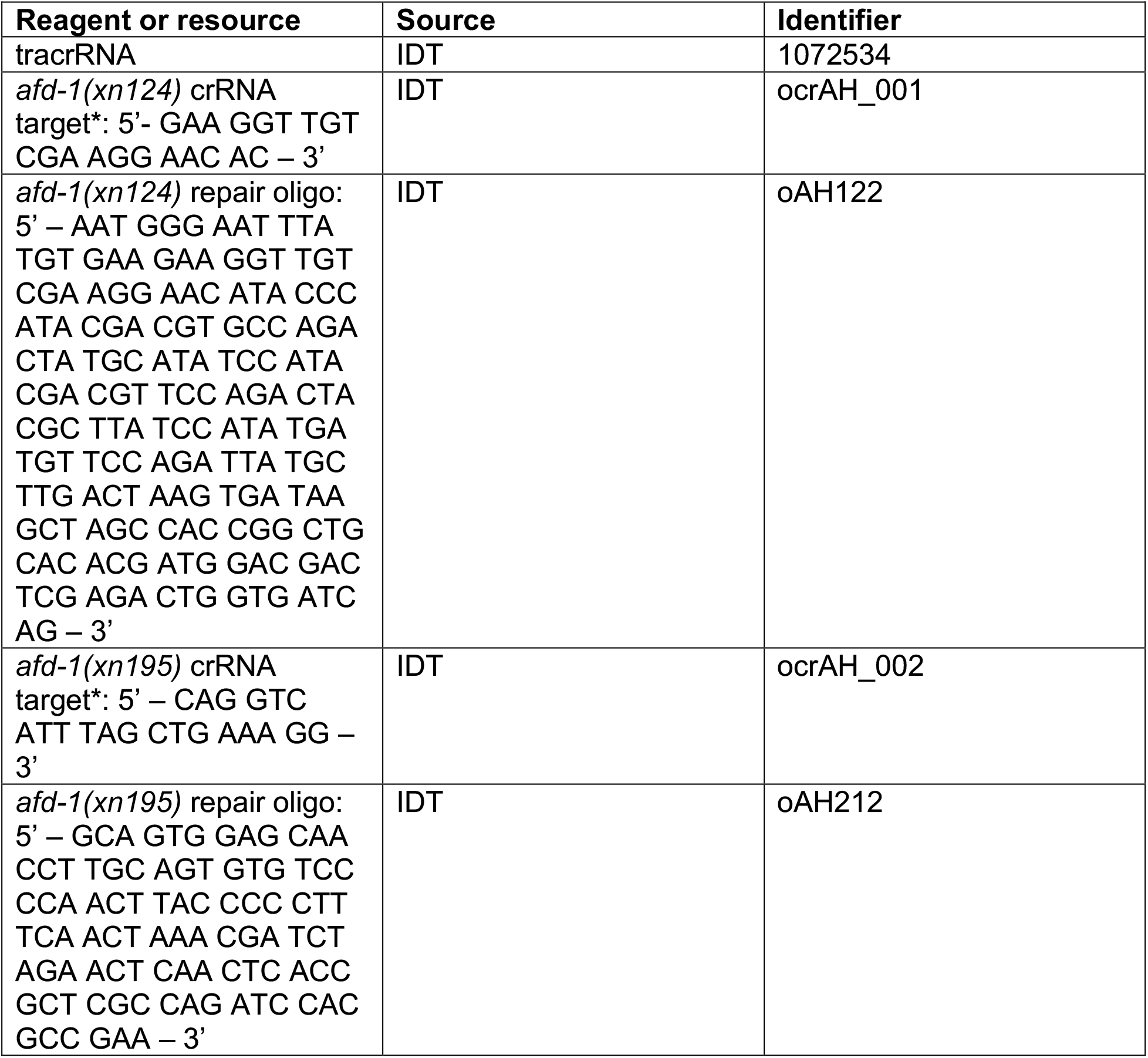
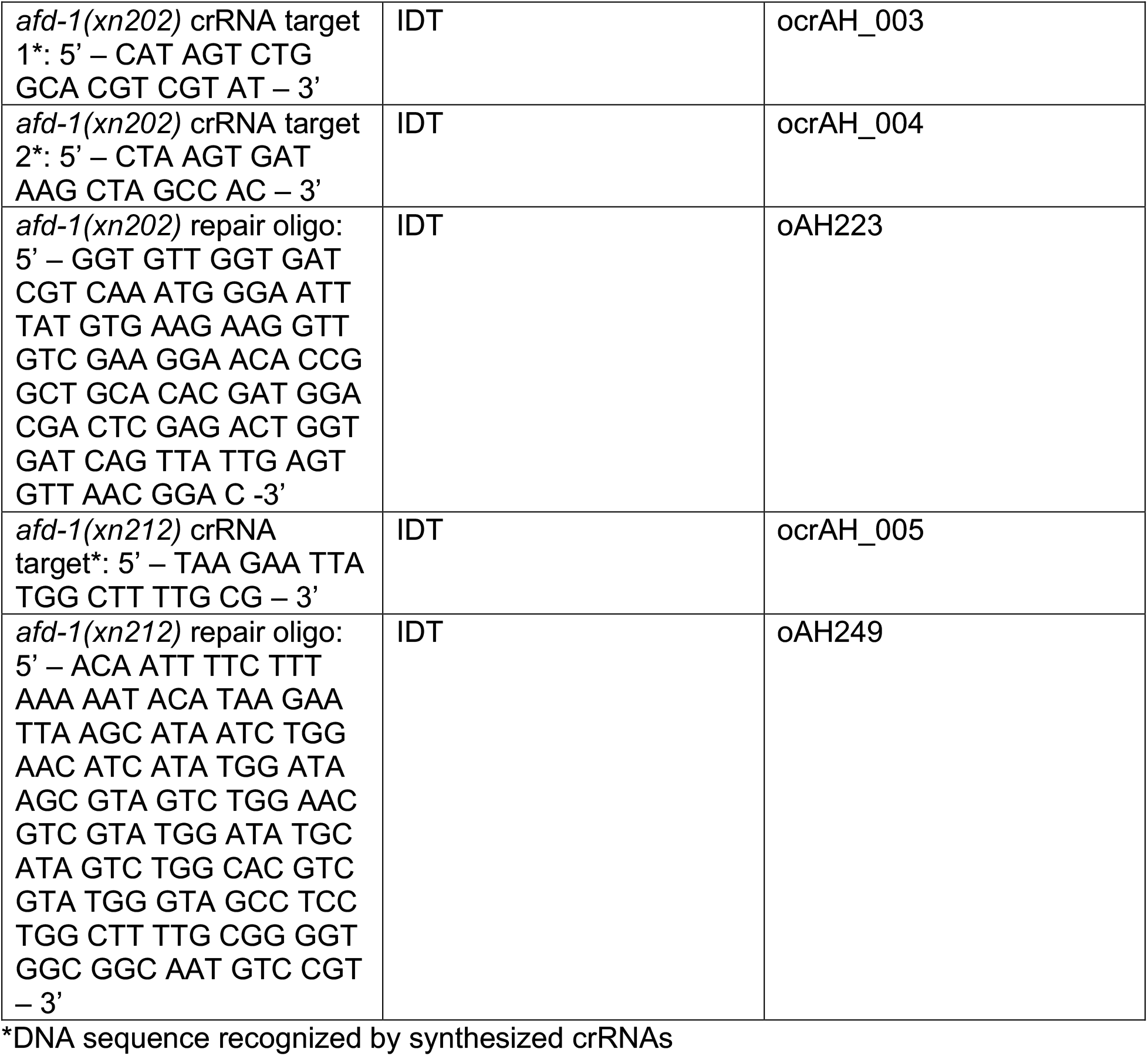

### Recombinant DNA

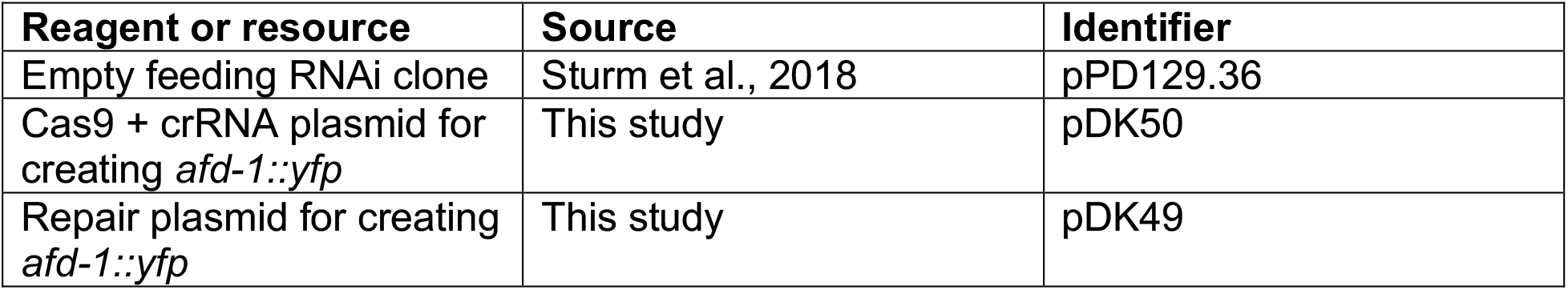

| Reagent or resource | Source | Identifier |
| --- | --- | --- |
| Empty feeding RNAi clone | Sturm et al., 2018 | pPD129.36 |
| Cas9 + crRNA plasmid for creating *afd-1::yfp* | This study | pDK50 |
| Repair plasmid for creating *afd-1::yfp* | This study | pDK49 |

